# MimicNeoAI: An integrated pipeline for identifying microbial epitopes and mimicry of tumor neoepitopes

**DOI:** 10.1101/2025.06.13.658292

**Authors:** Tao Chen, Wei Wang, Xiao Zuo, Yuxin Zhang, Mingwei Li, Zhilei Li, Yin He, Yanfei Zhou, Fang Ye, Bin Zhang, Qionghui Jiang, Huimin Liu, Lu Zhang, Jinman Fang, Yuanwei Zhang

**Author notes:** Corresponding author. (ZL); (FJM); (ZYW).

## Abstract

Tumor-associated microbial antigens represent promising immunotherapy targets, yet systematic identification methods remain underdeveloped. We developed MimicNeoAI, a computational pipeline integrating BiLSTM networks to identify microbial epitopes, mutation-derived neoepitopes, and their microbial mimics from sequencing data. Training on validated epitope datasets yielded 0.90 AUC with 91% accuracy on experimental validation sets. Application to colorectal cancer revealed that microbial epitopes, despite originating from a nine-fold smaller peptide pool, generated twice the immunogenic candidates (153 vs 75) compared to mutation-derived neoepitopes. These microbial epitopes exhibited exclusive tumor-specificity with no overlap in normal tissues. Single-cell TCR sequencing confirmed clonal expansion against 75% of predicted highly immunogenic epitopes, with molecular dynamics simulations demonstrating positive correlation between predicted immunogenicity and HLA-epitope-TCR binding stability. Collectively, our pipeline systematically unveils abundant, tumor-specific, and highly immunogenic microbial epitopes, providing a computational framework for developing broadly applicable cancer immunotherapies that leverage the tumor microbiome as an untapped source of therapeutic targets.

## Introduction

Cancer immunotherapy efficacy depends on identifying tumor-specific antigens that elicit robust T cell responses^1–3^. While mutation-derived neoantigens show promise, their extreme inter-patient heterogeneity limits clinical application to personalized therapies^4, 5^, necessitating alternative broadly applicable targets. Tumor-resident microbiomes represent an untapped immunogenic reservoir^6, 7^. Unlike patient-specific neoantigens, microbial antigens exhibit consistent enrichment patterns across patients, enabling broadly effective immunotherapies. Furthermore, microbial epitopes can trigger anti-tumor immunity through molecular mimicry—sharing structural similarities with tumor neoantigens while potentially exhibiting superior immunogenicity^8–10^. However, systematic exploration remains limited by computational constraints.

Current epitope prediction operates in silos: mutation-derived neoantigen tools cannot evaluate microbial peptides, while microbial epitope discovery lacks integration with neoantigen analysis ^5, 11–18^. Additionally, microbial epitope discovery confronts unique challenge: extracting low-abundance microbial sequences from host-dominated datasets. Consequently, no existing framework addresses these requirements within a unified analytical pipeline.

Accurate immunogenicity prediction remains a critical challenge, particularly for microbial epitopes. Although numerous algorithms effectively predict peptide-HLA binding^15, 19–21^, binding probability poorly correlates with immunogenicity—many high-affinity peptides fail to elicit T cell responses^22, 23^. This disconnect is amplified for microbial epitopes: existing immunogenicity prediction models^6, 24^, predominantly trained on mutation-derived neoepitope datasets, fail to capture microbial-specific immunogenic features. Moreover, current tools suffer from technical constraints—limited HLA allele coverage and restricted peptide length support—that hinder clinical translation. Deep learning architectures from natural language processing offer a transformative approach^25–27^. These architectures are well-suited for learning complex patterns across diverse peptide sources, including those crucial for identifying molecular mimicry. This dual-source analysis capability, essential for leveraging mimicry-based anti-tumor immunity, remains absent from existing platforms.

We developed MimicNeoAI, an integrated computational pipeline that simultaneously identifies tumor-associated microbial epitope candidates from whole-transcriptome sequencing (WTS) data and tumor mutation-derived neoepitope candidates from whole-exome sequencing (WES) data. MimicNeoAI employs a Bidirectional Long Short-Term Memory (BiLSTM)^27^ trained on validated epitopes from both microbial and host sources (IEDB and MicroEpitope^22,28^), incorporating peptide physicochemical properties and amino acid composition features. The model achieves comprehensive HLA coverage through direct integration of full-length sequences from IPD-IMGT/HLA^29^ and supports extensive peptide lengths. Crucially, MimicNeoAI identifies microbial epitopes exhibiting molecular mimicry with tumor mutation-derived neoepitopes.

Benchmarking demonstrates MimicNeoAI’s superior performance, particularly for microbial epitope candidates. Applied to colorectal cancer (CRC), microbial epitopes —despite originating from a nine-fold smaller peptide pool—yielded five-fold more high-immunogenicity candidates than mutation-derived counterparts. Single-cell T cell receptor sequencing (scTCR-seq) confirmed preferential T cell activation against predicted high-immunogenicity epitopes, with molecular dynamics simulations validating HLA-epitope-TCR binding stability. Thus, MimicNeoAI establishes a comprehensive framework for discovering broadly applicable cancer immunotherapy targets beyond patient-specific neoantigens.

## Results

### Overview of the MimicNeoAI pipeline

MimicNeoAI integrates parallel analytical pipelines to identify three epitope categories: tumor-associated microbial epitope candidates, mutation-derived neoepitope candidates, and their molecular mimic (Fig. 1). The microbial pipeline processes WTS data through host removing, taxonomic quantification, and peptide extraction, while the mutation-derived pipeline analyzes WES data via somatic variant calling and mutant peptide identification. Both pipelines converge on feature extraction, incorporating HLA typing and peptide physicochemical properties.

**Fig. 1.**
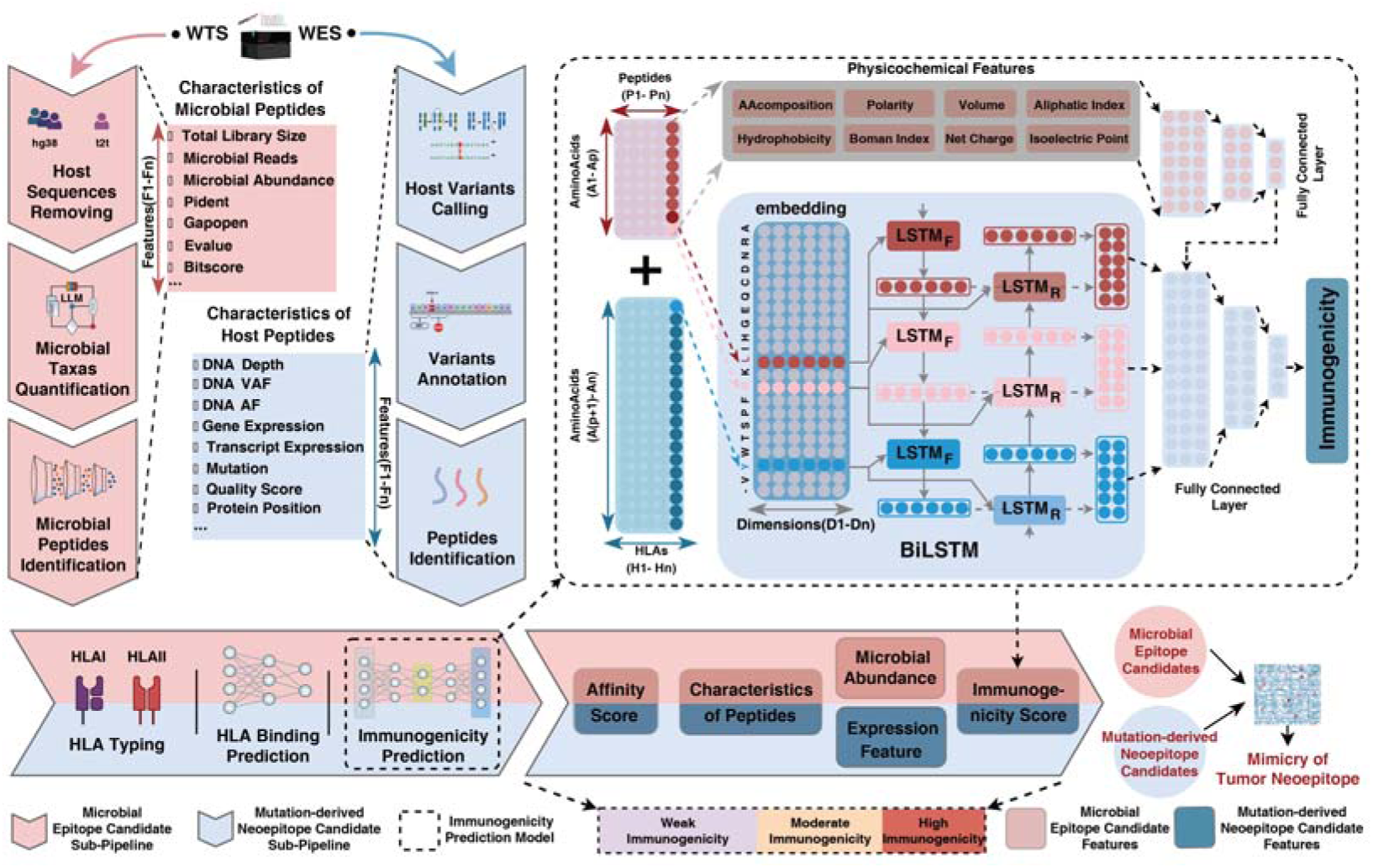
Overview of the MimicNeoAI pipeline. MimicNeoAI including two sub-pipelines for identifying microbial epitope candidates, mutation-derived neoepitope candidates, and their mimics of tumor neoepitope using WES and WTS data. For microbial epitope candidate identification, the pipeline executes host sequences removal, microbial taxa quantification, microbial peptides identification, microbial peptide characteristic generation, HLA typing, and immunogenicity prediction. For mutation-derived neoepitope candidate identification, the pipeline executes host variants calling, variants annotation, host peptide identification, peptide characteristic generation, HLA typing, and immunogenicity prediction. Both sub-pipelines employ an immunogenicity prediction model based on a combination of BiLSTM and FCNN. MimicNeoAI subsequently stratifies microbial epitope candidates and mutation-derived neoepitope candidates into distinct tiers based on HLA binding probability scores, immunogenicity scores, peptide characteristics, microbial abundance, and expression (detailed in Methods). Finally, mimicries of tumor neoantigen candidates are identified by assessing sequence similarity between microbial epitope candidates and mutation-derived neoepitope candidates.

Central to MimicNeoAI is a unified BiLSTM-based immunogenicity predictor that captures sequence-contextual patterns across both epitope sources. The network generates composite immunogenicity scores integrating HLA-binding probability, peptide characteristics, microbial abundance, and tumor gene expression, enabling stratification into distinct immunogenicity tiers (Supplementary Table 1). This dual-source architecture uniquely enables molecular mimicry detection through systematic sequence similarity evaluation between microbial and mutation-derived epitopes (detailed in Methods). Thus, MimicNeoAI transforms heterogeneous sequencing data into a prioritized catalog of immunotherapeutic targets spanning both microbial and mutational landscapes.

### Evaluation of the immunogenicity prediction performance

MimicNeoAI’s predictive performance was validated through computational benchmarking, experimental validation, ablation studies, and comparative analysis against state-of-the-art tools. Five-fold cross-validation on host-derived and microbial epitope datasets yielded a mean area under the receiver operating characteristic curve (ROC-AUC) of 0.909 ± 0.006 and a mean area under the precision-recall curve (PR-AUC) of 0.908 ± 0.007 (Fig. 2a, b; Supplementary Table 2). Converging training and validation losses confirmed model stability (Supplementary Fig. 1a-e). Experimental validation using 21 immunogenic and 21 non-immunogenic microbial peptide-HLA complexes demonstrated (19/21 true positives), 81.0% specificity (17/21 true negatives), accuracy of 0.857, precision of 0.826, recall of 0.905, and an F1 score of 0.864 (Fig. 2c, d; Supplementary Information; Supplementary Table 2). To investigate the impact of training data, we compared models trained exclusively on different epitope datasets: exclusively on microbial epitopes, exclusively on mutation-derived epitopes, and the combined dataset. Models trained exclusively on microbial epitopes outperformed those trained on mutation-derived epitopes for microbial prediction tasks (Fig. 2e, f; Supplementary Fig. 1f, g), suggesting microbial-specific biophysical properties. Nevertheless, the combined-dataset model maintained robust performance across both epitope types and was retained for comprehensive predictive capability (Fig. 2a, b).

**Fig. 2.**
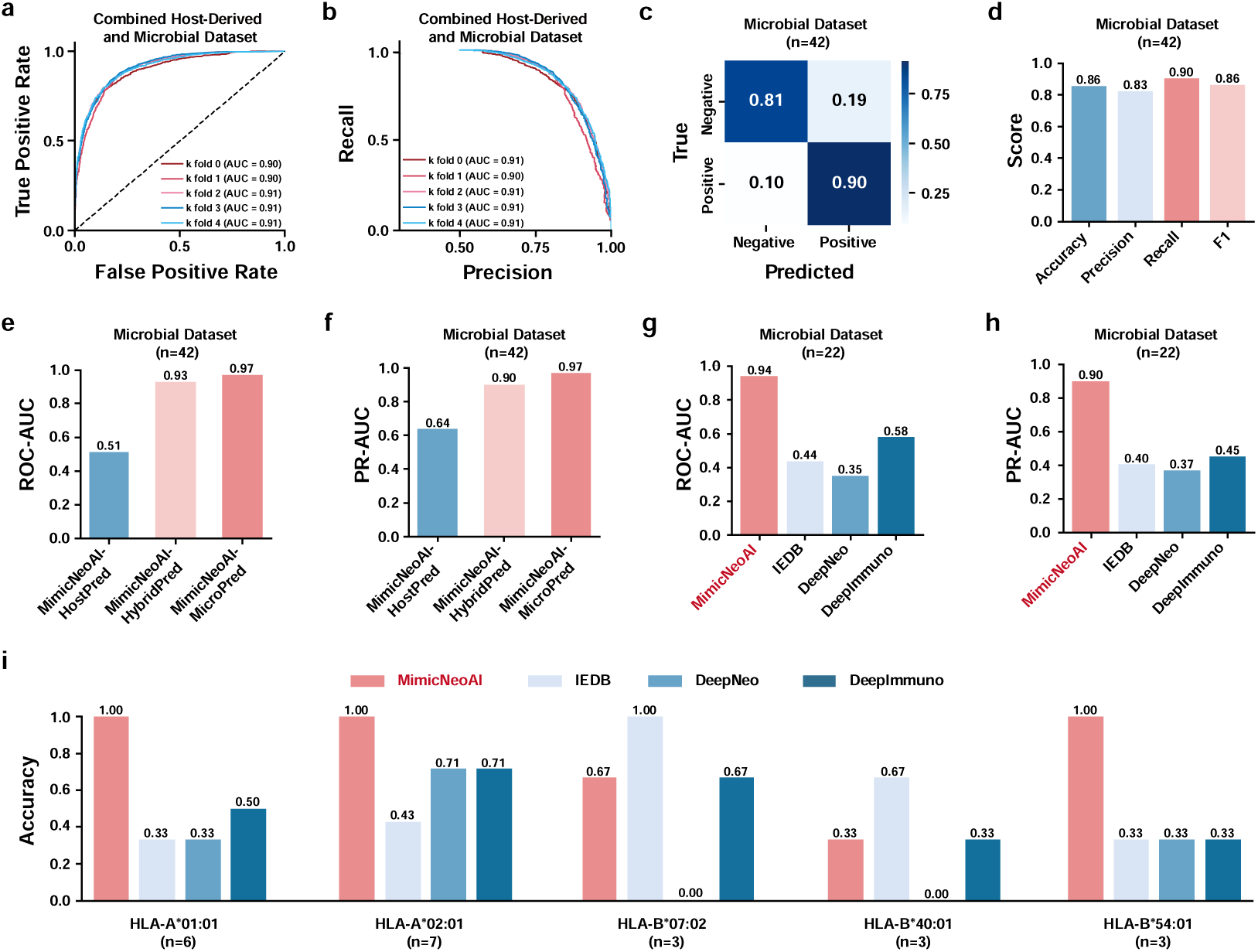
Performance evaluation of MimicNeoAI’s immunogenicity prediction model. (**a-b**) ROC (**a**) and PR curve (**b**) illustrating performance metrics based on five-fold cross-validation with datasets containing both host-derived epitopes and microbial epitopes. (**c-d**) Confusion matrix (**c**) and detailed performance metrics (**d**) showing precision using an additional validation dataset comprising 21 experimentally immunogenic microbial epitope-HLA complexes paired with 21 non-immunogenic controls. (**e-f**) Comparison of the ROC-AUC (**e**) and PR-AUC (**f**) across MimicNeoAI models trained on different epitope datasets (MimicNeoAI-HostPred: training host-derived epitopes only; MimicNeoAI-MicroPred: training with microbial epitopes only; MimicNeoAI-HybridPred: training with both host-derived epitopes and microbial epitopes). (**g-h**) Comparison of ROC-AUC (**g**) and PR-AUC (**h**) metrics between MimicNeoAI and established prediction algorithms (IEDB, DeepNeo, DeepImmuno), using a validation dataset comprising experimentally validated microbial antigens (22 microbial antigens supported by all algorithms). (**i**) Comparison accuracy of MimicNeoAI, IEDB, DeepNeo, and DeepImmuno on microbial epitopes presented by each HLA types (HLA-A*01:01, HLA-A*02:01, HLA-B*07:02, HLA-B*40:01, and HLA-B*54:01).

Comparative benchmarking against (IEDB^22^, DeepNeo^30^, DeepImmuno^31^) demonstrated MimicNeoAI’s superiority. On a 22 microbial epitope-HLA complexes compatible with all tools’ constraints (i.e., specific peptide lengths and HLA alleles supported by the competitors; see Supplementary Information for subset selection criteria), MimicNeoAI achieved ROC-AUC of 0.94 (vs. 0.35–0.58 for comparator tools, P = 2.567×10^−4^ vs IEDB, P = 5.607×10^−6^ vs DeepNeo, P = 1.326×10^−2^ vs

DeepImm, DeLong’s test) and a PR-AUC of 0.90 (vs. 0.37–0.45 ) (Fig. 2g, h; Supplementary Table 3). Critically, MimicNeoAI’s unrestricted peptide length and comprehensive HLA coverage enabled analysis of all experimental epitopes in the IPD-IMGT/HLA database, whereas comparator excluded 40.5%, 38.1%, and 0% of initial experimental epitopes in IEDB, DeepNeo, and DeepImm respectively due to algorithmic restrictions. Notably, within the 22-epitope subset, MimicNeoAI achieved 100% accuracy for epitopes associated with common HLA alleles (HLA-A*01:01, HLA-A*02:01, and HLA-B*54:01) within the comparison subset (Fig. 2i).

### Pilot Application of MimicNeoAI in a Colorectal Cancer Case Study

We applied MimicNeoAI to tumor and normal colorectal tissue on CRC, analyzing both WTS and WES data to demonstrate the complete analytical pipeline (Fig. 3a-d). Microbial profiling revealed substantial differences between the tumor and normal microbiomes in this patient. Taxa enriched in the tumor tissue were absent or diminished in normal tissue (Fig. 3a, b). Known CRC-associated species (*Bacteroides fragilis* and *Fusobacterium nucleatum*^32, 33^) co-occureed with supected CRC-linked species (*Prevotella copri* and *Bilophila wadsworthia*^34, 35^). Oral cavity-resident species (*Aggregatibacter aphrophilus* and *Aggregatibacter segnis*^36–39^) were identified in the tumor tissue, consistent with the hypothesis of bacterial translocation from the oral cavity.

**Fig. 3.**
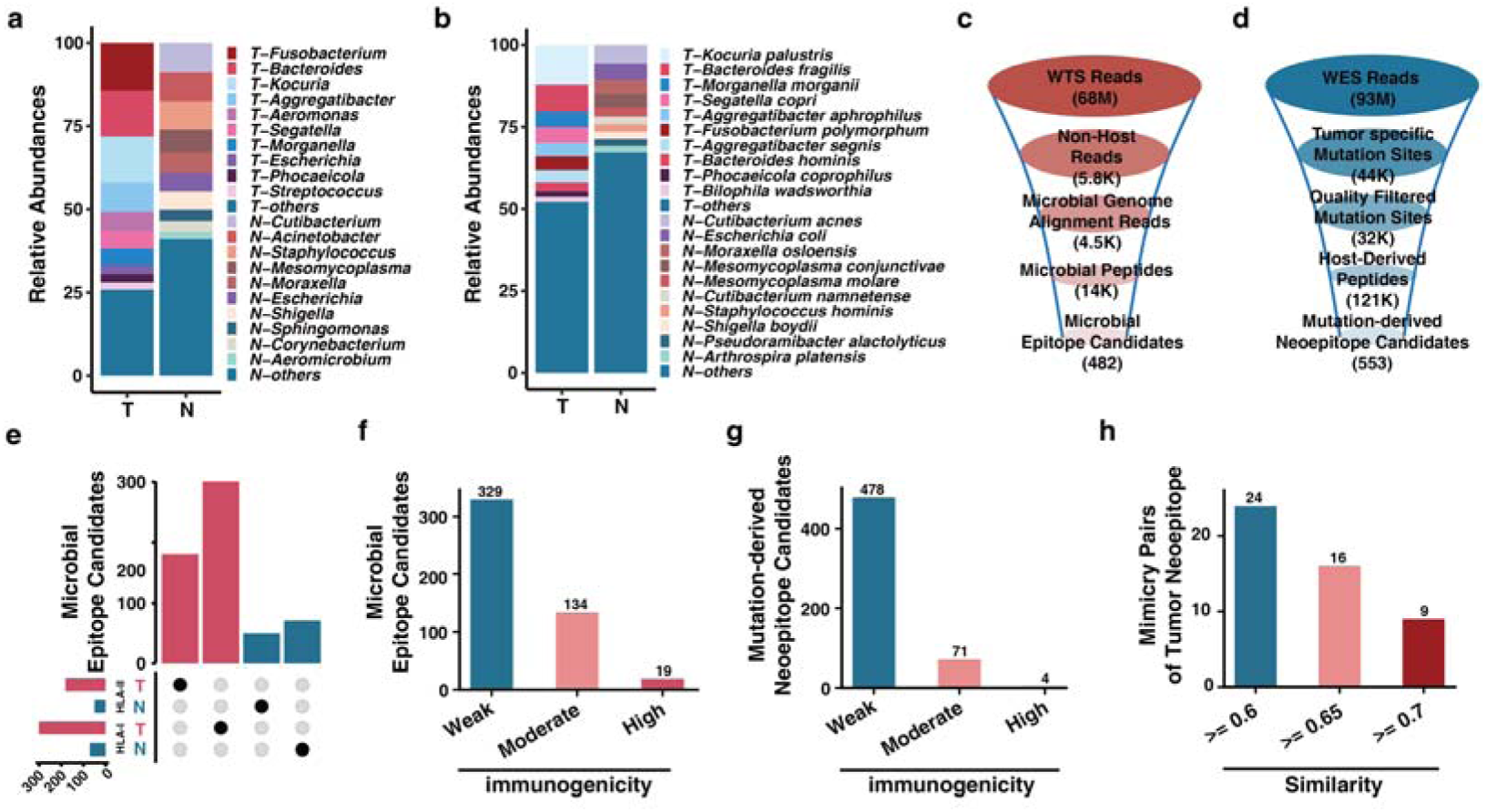
Identification of microbial epitope, mutation-derived neoepitope candidates and their mimics of tumor neoepitope in CRC. (**a-b**) Relative abundances of the top 10 microbial genera (**a**) and species (**b**) detected in tumor versus normal samples. (**c-d**) Data statistics summary of the number of microbial (**c**) and mutation-derived (**d**) epitope candidates identified by the MimicNeoAI pipeline. (**e**) UpSet plot showing intersections of microbial epitope candidates uniquely identified in tumor (red) or normal (blue) samples, separately presented by HLA-I and HLA-II. (**f-g**) Number of microbial (**f**) and mutation-derived (**g**) epitope candidates categorized according to immunogenicity tiers: weakly, moderately, and highly. (**h**) The number of tumor neoantigen mimicry pairs presented by HLA-I.

MimicNeoAI processed 68 million tumor WTS reads, extracting 5,782 high-quality non-host reads. Microbial genome alignment retained 4,500 reads, yielding 13,950 unique microbial peptides after in silico translation. HLA binding probability (HLA-I/II) and immunogenicity prediction identified 482 tumor-specific microbial epitope candidates (HLA-I: 301, HLA-II: 181; Fig. 3c), with no identical microbial epitopes identified in the normal colorectal tissue (Fig. 3e). Parallel WES analysis of 93 million reads identified 44,248 somatic mutations, with 32,405 high-confidence sites retained after quality filtering. These mutations generated 121,260 mutation-derived peptides, yielding 553 tumor-specific mutation-derived epitope candidates after HLA binding and immunogenicity prediction (HLA-I: 451, HLA-II: 102; Fig. 3d).

MimicNeoAI was used to categorize epitope candidates into three immunogenicity levels—weak, moderate, and high—based on thresholds tailored to the data source (see Methods for details). Microbial candidates yielded a total of 19 high, 134 moderate (a total of 153 in high-moderate group), and 329 weak immunogenicity microbial epitope candidates (Fig. 3f). Mutation-derived candidates produced 4 high, 71 moderate (a total of 75 in high-moderate candidates), and 478 weak immunogenicity epitope candidates (Fig. 3g, Supplementary Table 4). Despite originating from a 9-fold smaller peptide pool (13,950 vs 121,260), microbes generated twice the high-moderate immunogenicity (153 vs 75) and nearly five-fold more high immunogenicity epitope candidates (19 vs 4). Molecular mimicry analysis identified 24 HLA-I-restricted candidate pairs with similarity scores >0.6 (detailed in Methods), all sharing ≥6 consecutive identical amino acids. Nine pairs exhibited 7-residue identity stretches, suggesting potential cross-reactive immunity between microbial and mutation-derived epitopes (Fig. 3h; Supplementary Table 5).

### Assessment of Predicted Microbial Epitope Immunogenicity via scTCR-Seq

The scTCR-seq analysis of peripheral blood mononuclear cells (PBMCs) from a therapy-responsive CRC patient validated MimicNeoAI’s immunogenicity predictions through TCR clonal expansion patterns. Clonally expanded TCR clonotypes (>1 cell with identical CDR3 sequences), represented 8% of total clonotypes but comprised 60% of TCR read counts (Fig. 4a), indicating antigen-driven expansion. We then evaluated the binding probability between TCR and epitope against 301 HLA-I-restricted microbial epitope candidates (Supplementary Table 6, detailed in Method). The TCRs were stratified as: High probability (mean binding probability ≥0.6 and clonal expansion), Medium probability (0.4 ≤mean binding probability <0.6 and expanded), and Low probability (mean binding probability <0.4 or non-expanded) groups. Expanded clonotypes showed higher probability scores than non-expanded counterparts (Fig. 4b), with positive correlation between clonal expansion (read count ratio) and mean TCR binding probability (*r* = 0.50, P = 7.244 × 10□^30^; Fig. 4c).

**Fig. 4.**
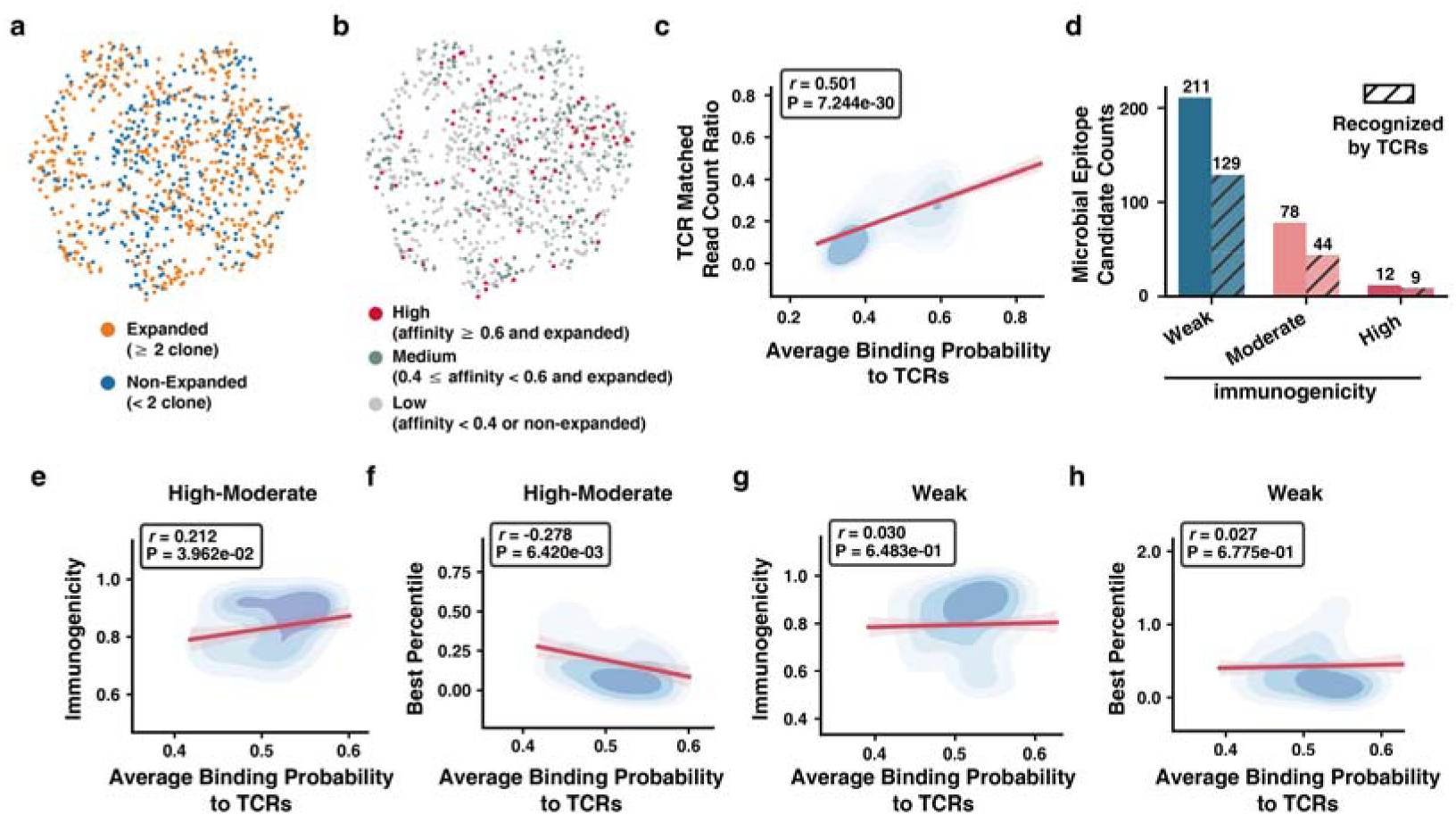
Validation of microbial epitope candidate immunogenicity via TCR binding probability and clonal expansion. (**a**) Clustering distribution of TCR categorized by clonal expansion status (expanded: ≥ 2 clone; non-expanded: < 2 clone). (**b**) Clustering distribution of TCR stratified by combined predicted binding probability and clonal expansion status: high (binding probability ≥ 0.6 and expanded), medium (0.4 ≤ binding probability < 0.6 and expanded), and low (binding probability < 0.4 or non-expanded). (**c**) The correlation between TCR read count ratios and binding probability. (**d**) Distribution of HLA-I presented and TCR recognized microbial epitope candidates across immunogenicity tiers (weak, moderate, and high). TCR recognized microbial epitope candidates indicated by diagonal-striped bars. (**e-f**) Correlation between immunogenicity score (**e**) and best percentile (**f**) and average clonally expanded TCR binding probability for microbial epitope candidates in High-Moderate group. (**g-h**) Correlation between immunogenicity score (**g**) and best percentile (**h**) and average clonally expanded TCR binding probability for microbial epitope candidates in Weak group.

We conducted a further assessment of the association between predicted immunogenicity and TCR clonal expansion capacity. Epitopes were classified as functionally immunogenic if: (i) Higher mean binding probability to expanded vs. non-expanded TCRs (P = 6.501 × 10□^34^, Mann–Whitney test), and (ii) Mean binding probability to expanded TCRs >0.5. High-immunogenicity epitopes were enriched among those preferentially recognized by expanded TCRs (75%, 9/12) versus moderate (56%, 44/78) and weak (61%, 129/211) epitopes (Fig. 4d). Read counts of expanded TCRs correlated positively with average binding probability across all immunogenicity tiers (*r* = 0.460 for High-Moderate, P = 7.210 × 10□^25^; *r* = 0.511 for Weak, P = 3.083 × 10□^31^; Supplementary Fig. 2a, b).

Next, we investigated the correlation between the predicted immunogenicity and TCR recognition by assessing associations between immunogenicity score, best percentile (detailed in Methods), and average probability to expanded TCRs. Within High-Moderate immunogenicity group, immunogenicity scores positively correlated with average binding probability to expanded TCRs (*r* = 0.212, P < 0.05; Fig. 4e), while best percentile showed negative correlation (*r* = -0.278, P = 6.420 × 10□³; Fig. 4f). No such correlations existed for weak epitopes (Fig. 4g, h). Consistent with established HLA-I binding principles^40–42^, amino acid frequency analysis confirmed that anchor positions P2 and P9 in the predicted 9-mer epitopes were predominantly occupied by hydrophobic residues (Supplementary Fig. 2c-f). MimicNeoAI effectively prioritizes epitopes with enhanced TCR engagement potential, particularly at higher immunogenicity tiers.

### Validation of TCR-epitope-HLA interactions using molecular dynamics simulations

Molecular dynamics simulations provided structural validation of MimicNeoAI’s predictions using representative HLA-peptide-TCR complexes from (i) *Bacteroides fragilis* (CRC-associated), (ii) HLA-A*02:01*, and (iii) the dominant T cell clonotype specific for *HLA-A*02:01-restricted 9-mer peptides with high TCR binding probability (≥0.8). Two high immunogenicity microbial epitope candidates (FLLCLVCVL, KLADAFTPL) and three weak immunogenicity candidates (MSYAAIPGV, SAFIVLFAV, YTNPEIAGV) were selected for analysis.

Molecular dynamics simulations revealed negative binding free energies (ΔG) for four of five complexes (Fig. 5a; Supplementary Table 7). High immunogenicity microbial epitope candidates exhibited substantially stronger binding (mean ΔG = -14.46 kcal/mol) than weak immunogenicity counterparts (mean ΔG = -1.28 kcal/mol). Structural analysis of the most stable complex highlighted key binding interfaces (Fig. 5b), while full structural ensembles shown for all complexes (Fig. 5c-f). Production trajectories at 0, 50, and 100 ns confirmed structural equilibrium throughout simulations (Fig. 5g; Supplementary Fig. 3a-d).

**Fig. 5.**
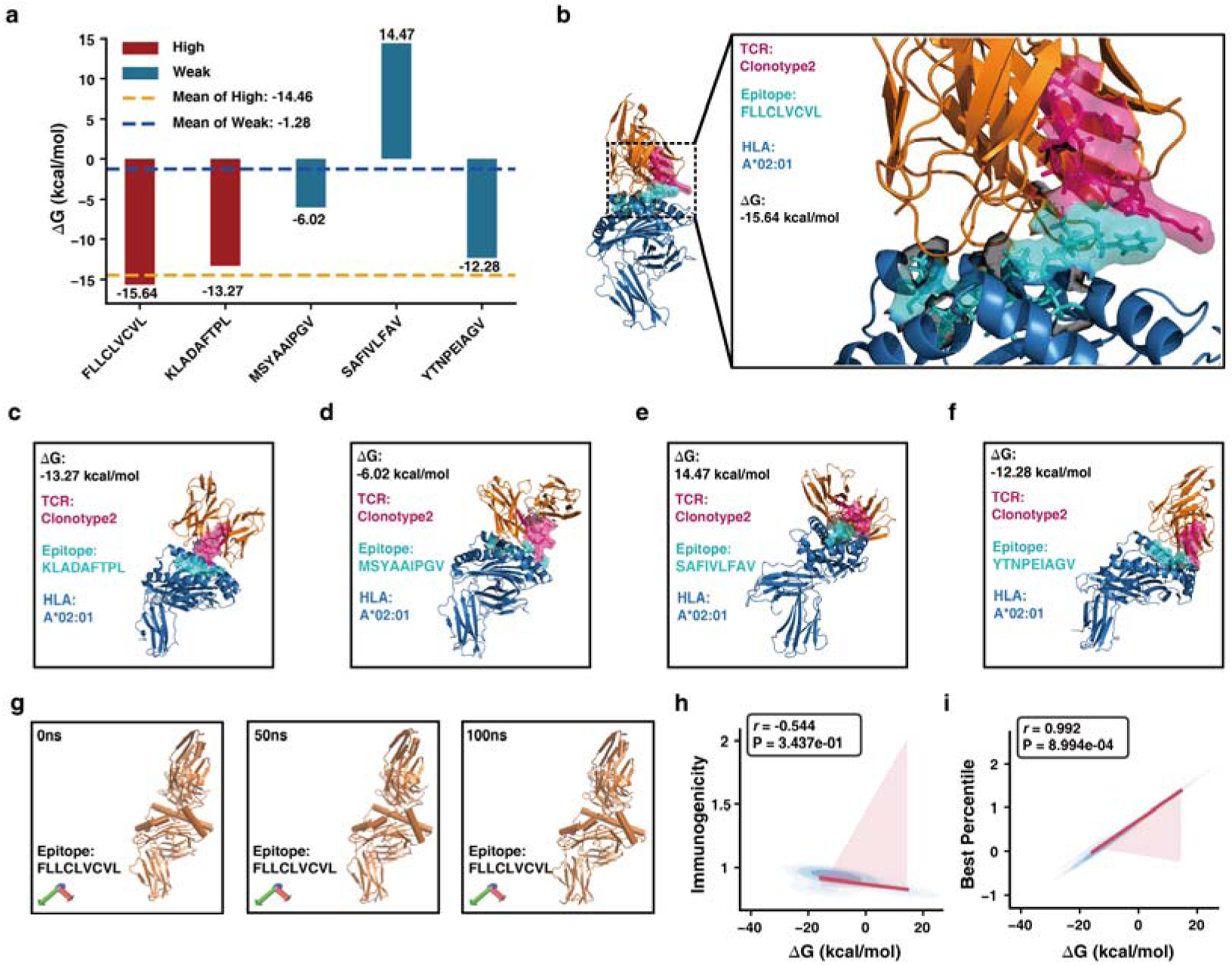
Assessment of TCR-epitope-HLA interactions via molecular dynamics simulations. (**a)** The ΔG of microbial epitope candidates with high and weak immunogenicity. (**b**) Structural representation of the interaction among Clonotype2, HLA-A*02:01, and epitope FLLCLVCVL. (**c-f**) Structural representations of Clonotype2 and HLA-A*02:01 complexed with the microbial epitope candidates: KLADAFTPL (**c**), MSYAAIPGV (**d**), SAFIVLFAV (**e**), and YTNPEIAGV (**f**). (**g**) The production simulation trajectory over 100 ns is divided into three time interval (0 ns, 50 ns, and 100 ns). (**h-i**) Correlation analyses of ΔG with immunogenicity scores (**h**) and best percentiles (**i**) for microbial epitope candidates.

Pearson correlation analysis revealed a negative correlation between immunogenicity score and ΔG (*r* = -0.544, P = 3.437×10^−1^; Fig. 5h), while best percentile showed near-perfect positive correlation (*r* = 0.992, P = 8.994×10^−4^; Fig. 5i). Additional metrics, including best IC50 score (*r* = 0.828, P = 8.344×10^−2^), median percentile (*r* = 0.581, P = 7449×10^−1^), and median IC50 score (*r* = 0.202, P = 3.045×10^−1^) exhibited consistent trends (Supplementary Fig. 3e-g). Thus, molecular dynamic simulations confirms that MimicNeoAI-predicted immunogenicity corresponds to energetically stable ternary complexes, while superior HLA binding predictions associate with favorable interaction thermodynamics.

## Discussion

Cancer immunotherapy requires broadly applicable target antigens beyond patient-specific neoantigens. MimicNeoAI addresses this need through unified computational discovery of microbial epitope and mutation-derived neoepitope candidates, revealing tumor-associated microbes as an untapped source of shared immunotherapy targets.

MimicNeoAI’s integration of microbial and host genomic analysis within a single pipeline enables comprehensive epitope candidate discovery from WTS and WES data. Benchmarking demonstrated competitive performance in immunogenicity prediction (ROC-AUC >0.90 across datasets), with notably improved accuracy for microbial epitopes versus IEDB, DeepNeo, and DeepImmuno (Fig. 2d-e; Supplementary Table 3). This enhancement stems from our innovative approach of training on host epitope datasets with the additional inclusion of microbial epitope datasets, coupled with the integration of peptide physicochemical properties and BiLSTM-based sequence feature extraction. Unrestricted peptide length support and comprehensive HLA coverage further distinguish MimicNeoAI from current methods constrained by algorithmic limitations.

Application to CRC revealed striking disparities between epitope sources. Despite originating from a nine-fold smaller peptide pool, MimicNeoAI identified comparable numbers of microbial epitope (n = 482) and mutation-derived neoepitope (n = 553) candidates. Microbial candidates yielded twice the high-moderate immunogenicity epitopes compared to mutation-derived neoepitope candidates (153 vs 75), suggesting that microbes may contribute to stronger immunogenic potential in this tumor context. The complete non-overlap microbial epitope candidate between tumor and normal colorectal tissue (Fig. 3e) confirmed tumor-specificity. Detection of CRC-associated bacteria (*Bacteroides fragilis* and *Fusobacterium polymorphum*) validated our microbial profiling approach.

The scTCR-seq analysis demonstrated that 75% of predicted high immunogenicity epitope candidates were recognized by clonally expanded TCRs (Fig. 4d), with immunogenicity scores correlating with TCR reactivity for high-moderate candidates (Fig. 4f). Molecular dynamics simulations revealed stronger binding energetics for high immunogenicity complexes (mean ΔG = -14.46 vs -1.28 kcal/mol, Fig. 5a, h), with near-perfect correlation between predicted HLA binding and thermodynamic stability (*r* = 0.992, P = 8.994×10□□; Fig. 5i).

Current limitations include single-patient validation requiring multi-center cohort expansion, potential enhancement through structural feature integration (e.g., AlphaFold2 embeddings), and need for systematic biological validation via high-throughput TCR binding assays. Nevertheless, MimicNeoAI establishes a computational framework for leveraging the tumor microbiome as a prolific source of immunotherapy targets, potentially transforming cancer treatment from personalized to broadly applicable strategies.

## Methods

### In-house sequencing data generation

Tumor tissue, adjacent normal tissue, and PBMCs were obtained from CRC patients. Tumor specimens were obtained via endoscopic biopsy, with approximately 2–3 pieces (∼3 mm in diameter each) excised based on tumor dimensions. Samples were immediately rinsed with saline solution to eliminate residual blood and mucus, snap-frozen in dry ice for approximately 2 hours during transportation, and subsequently stored at −80 °C until further processing. PBMCs were isolated from the patient’s peripheral blood using density gradient centrifugation. Blood samples were mixed with an equal volume of phosphate-buffered saline (PBS), followed by centrifuged at 800 × g for 20 minutes. The PBMC fraction was collected, washed twice with PBS, resuspended in cryopreservation medium, and stored at −80 °C.

For WES, DNA was extracted using the HiPure Universal DNA Kit (D3018-02, Magen). DNA sequencing libraries were constructed using the Hieff NGS® OnePot Pro DNA Library Prep Kit for Illumina® (12205, Yeasen), and subsequently sequenced on the DNBSEQ-T7 sequencing platform, generating 150 bp paired-end reads.

For WTS, total RNA was isolated using the Hieff NGS® MaxUp Human rRNA Depletion Kit (12257ES24, Yeasen). RNA quality and integrity were verified with the Agilent 2100 Bioanalyzer. Transcriptome libraries were prepared utilizing the Hieff NGS® Ultima Dual-mode mRNA Library Prep Kit (12301ES96, Yeasen), followed by sequencing on the DNBSEQ-T7 platform.

For scTCR-seq, PBMCs were isolated by density gradient centrifugation using Ficoll-Paque plus (04-03-9391/03, Stemcell). Single-cell sequencing libraries were generated using the Chromium Single Cell 5’ v2 and V(D)J Reagent kits (10x Genomics). Library quality and concentration were assessed by an Agilent Bioanalyzer 2100 with the High Sensitivity DNA Chip and a Qubit High Sensitivity DNA Assay (Thermo Fisher Scientific). Sequencing was performed on the DNBSEQ-T7 platform, producing 150-bp paired-end reads.

### Host sequences removal

Host genomic sequences were removed using a two-step alignment strategy against the GRCh38^43^ and T2T-CHM13v2.0^44^ reference genomes. Quality-controlled sequencing reads (processed with Fastp^45^ v0.22.0) were initially aligned to GRCh38 using BWA-MEM^46^ (v0.7.17). Unmapped reads were extracted with SAMtools^47^ (v1.5), converted to FASTQ format, and realigned to T2T-CHM13v2.0 with BWA-MEM. eads remaining unaligned to both reference genomes were considered non-host sequences. Alignment metrics (total reads, mapped reads and paired-end reads) were documented at each alignment step using SAMtools. For paired-end sequencing data, unmapped reads are separated into Rl and R2 FASTQ files for each sample; for single-end sequencing data, unmapped reads were consolidated into one FASTQ file per sample.

### Custom microbial reference database construction

Taxonomic data relevant to human tumor was initially curated from the Human Microbiome Project (HMP^48^) , published reference genomes [Thomas et al.]^49^, and literature. We identified 355 bacterial genera and established their taxonomic hierarchy using NCBI taxonomy ID (ftp://ftp.ncbi.nlm.nih.gov/pub/taxonomy/taxdump.tar.gz, updated on Nov. 7, 2024.).

For these genera, 8,315 high-quality reference genomes (species or strain level) were retrieved from NCBI RefSeq using assembly summary data (ftp://ftp.ncbi.nlm.nih.gov/genomes/refseq/assembly_summary_refseq.txt, updated on Nov. 14, 2 024.). These genomes, manually curated by the NCBI, met high-quality standards. Viral reference sequences were sourced from Thomas et al^49^. and NCBI Re fSeq.

To assess colonization potential, tumorigenic associations, and effects on anti-tumor treatments, we employed a large language model (LLM, ChatGPT 4.0) with the prompt:

“The list below contains microbiome taxa at the species or strain level. Please provide the following information for each and organize it in a table: Brief Description, Potential to Colonize Human Tissues, Association with Tumorigenesis and Progression, and Impact on Anti-Tumor Treatments.”

Taxa were queried in randomized batches of 50. LLM outputs underwent manual validation via literature review and expert curation, retaining 3,199 bacterial and 1,262 viral taxa. Criteria included:

Colonization Potential: Low (minimal tissue presence), Moderate (tissue-specific, non-dominant), High (prevalent in tissues).

Tumorigenic Association: Established (e.g., Porphyromonas gingivalis), Opportunistic (e.g., Enterobacter asburiae), or Probiotic-linked (limited evidence; e.g., Limosilactobacillus panis).

Treatment Impact: Positive (e.g., Streptomyces spp. as drug sources), neutral, or negative (e.g., opportunistic pathogens complicating therapy).

Genome sequences were downloaded from NCBI RefSeq (ftp://ftp.ncbi.nlm.nih.gov/refseq/release/bacteria/*.fna.gz, ftp://ftp.ncbi.nlm.nih.gov/refseq/release/viral/viral. 1.1.genomic.fna.gz, updated on Sep. 7, 2024.). A reference database was built using BWA-MEM and GATK^50^ PathSeqBuildReferenceTaxonomy (v4.6). Protein sequences retrieved from NCBI RefSeq (ftp://ftp.ncbi.nlm.nih.gov/genomes/all/, updated on Sep. 7, 2024) and formatted with BLAST^51^ makeblastdb (v2.14.0).

### Microbial sequence alignment

Host-depleted sequencing reads were aligned to the custom microbial database using GATK PathSeqPipelineSpark^52^. Alignment parameters included retention of primary and suboptimal alignments and a minimum clipped read length of 50 bp. Taxonomic assignments were validated against a curated hierarchy file. Ambiguous alignments were resolved by discarding reads with alignment lengths less than 95% of the best match length, and excluding suboptimal alignments containing more than one mismatch.

### Microbial peptide identification

Taxon-filtered reads were queried against the custom protein database using BLASTX (v2.12.0) (E-value ≤ 5e-2, top 5 hits per query (-max_target_seqs 5)). Only alignments with 100% sequence identity to reference proteins (Pident = 100) were retained. Output fields included: qseqid, sseqid, qseq, sseq, stitle, pident, length, mismatch, gapopen, evalue, and bitscore. Validated peptides were converted to

FASTA format, and taxonomic origins were confirmed via NCBI taxonomy IDs. The FASTA files were then input into PVCTools to generate peptide sequences with lengths of 8-10 for HLA-I and 15 for HLA-II.

### Host variants identification, annotation and peptide identification

Raw sequencing data in FASTQ format were quality control using Fastp to remove adapter sequences and low-quality reads. Filtered reads were aligned to the human reference genome (GRCh38) using BWA-MEM with multiple threads enabled (parameter -t). SAM alignment files were converted to sorted BAM format and indexed using SAMtools with parallel sorting (-@ threads). Duplicate reads were identified and marked with GATK Spark (MarkDuplicatesSpark) with local multithreading (--spark-master local[threads]).

Base quality score recalibration (BQSR) was performed in two sequential steps. First, recalibration models were generated using GATK Spark BaseRecalibratorSpark, incorporating variant datasets (Mills_and_1000G_gold_standard.indels.hg38.vcf.gz, dbsnp_138.hg38.vcf.gz, and 1000G_phase1.snps.high_confidence.hg38.vcf.gz). Second, adjustments were applied with ApplyBQSRSpark. Somatic variant calling was subsequently conducted on BQSR-processed BAM files using GATK Mutect2, and candidate mutations were filtered using FilterMutectCalls to enhance specificity and reliability. Identified variants were annotated using VEP^53^ (v110) within Singularity^54^ containers (v3.9.0-rc.3). Annotation leveraged the human reference genome GRCh38 with the VEP cache and genome reference data mounted via the -B flag. Key parameters included: --offline mode with local cache, --vcf output format, transcript annotations (--symbol, --hgvs, --tsl, --biotype), and plugins (Frameshift, Wildtype) from /vep_data/VEP_plugins-release-110/. Population allele frequencies (--af, --af_1kg) and SIFT predictions (--sift b) were incorporated. Post-annotation filtering removed transcript-reference mismatches using ref-transcript-mismatch-reporter with strict mode (-f hard). The VCF file was then input into PVCTools to generate peptide sequences with lengths of 8-10 for HLA-I and 15 for HLA-II.

### HLA typing identification

HLA alleles were determined using HLA-HD^55^ (v1.7.0). The human HLA Class I loci (A, B, C, E, F, G) and Class II loci (DRA, DRB1, DQA1, DQB1, DPA1, DPB1) were assessed. FASTQ files were aligned to HLA references via Bowtie2^56^ (v2.2.8; parameters: -x [index] -p [threads]). Mapped reads were extracted (SAMtools view -h -F 4), converted to FASTQ, and standardized using AWK. HLA allele typing was performed using hlahd.sh (parameters: -m 50, population frequency data -f, HLA gene references).

### HLA binding prediction

Peptide-HLAs binding prediction were analyzed using pVACtools (v4.2.1)^18, 57, 58^. Epitope lengths: 8–10-mers (Class I), 15-mers (Class II). Prediction algorithms: NNalign, NetMHCpan, NetMHCIIpan, NetMHCpanEL, NetMHCIIpanEL, SMM, SMMPMBEC, and SMMalign. Predication results: Best IC50 Score, Best IC50 Score algorithm, Best Percentile, Best Percentile algorithm, Median IC50 Score, and Median Percentile, detailed scription in Supplementary Table 1.

### Immunogenicity prediction model construction

A Bidirectional LSTM (BiLSTM) ensemble model predicted immunogenicity for peptide-HLA pairs. Input encoding included:

Sequence embedding: Peptide and HLA sequences were tokenized using a 22-token vocabulary comprising a padding token (“<PAD>”), the 20 standard amino acid residues, and “-” representing the ambiguous amino acid X. The padding token was mapped to a 12-dimensional zero vector. The 20 standard residues and “-” were mapped to 12-dimensional vectors through a simplified PCA feature space derived from 566 physicochemical properties in the AAindex1 database^31, 59^.

Physicochemical features: Eight peptide-specific properties (Amino acid composition, polarity, volume, charge, hydrophobicity, boman index, aliphatic index, and isoelectric point) were computed using the R Peptides^60^ package.

Features were processed through a 256-node dense layer. HLA sequences (retrieved from IPD-IMGT/HLA^29, 61^ database) were aligned to HLA full-length amino acid sequences. Outputs were calibrated via softmax activation, and the probability value at index 1 was taken to represent the predicted probability of the positive class (immunogenic). Alleles without HLA full-length amino acid sequence matches received a neutral score (0.5).

### Model training and performance evaluation

The model parameters were optimized using the Adam optimizer (learning rate: 5×10□□) with a cross-entropy loss. A 5-fold cross-validation was employed, where each fold was trained for 200 epochs, requiring approximately 20-30 minutes of computation time per fold on a 48GB NVIDIA L20 GPU. In addition to cross-validation, formal model training was conducted using all data from the 5-fold cross-validation. All implementations leveraged GPU acceleration with jupyterlab-pytorch:2.0.1-py3.9-cuda11.7-ubuntu20.04 for improved efficiency.

### Microbial epitope candidate taxonomic resolution and annotation

To resolve taxonomic ambiguities, peptide-protein matches were cross-referenced against microbial taxonomic hierarchies and annotated protein databases. Peptides were retained only if they exhibited species-level consistency between aligned sequences and validated microbial taxon IDs. This ensured unambiguous linkage of microbial epitope candidates to specific species through both sequence alignment and taxonomic lineage validation.

Comprehensive features and annotations were integrated to support epitope evaluation, including: (i) immunogenicity classification (predicted immunogenicity tier by MimicNeoAI); (ii) peptide characteristics (sequence, length, and extended sequence information); (iii) HLA presentation (predicted HLA allele); and (iv) nucleotide- and taxon-level annotations (NCBI taxonomy ID, reference genome ID, scientific name, read counts, abundance metrics such as normalized abundance). Additionally, protein-level information was incorporated, comprising matched protein names and identifiers, as well as alignment quality metrics (bitscore, pident, mismatch, gapopen, and E-value). Binding probability annotations include the best and median IC50 values and percentile ranks across multiple algorithms, alongside the predictive method for each. Furthermore, immunogenicity scores predicted by MimicNeoAI and a series of physicochemical properties—such as amino acid composition, polarity, volume, net charge, hydrophobicity, Boman index, aliphatic index, and isoelectric point—are also provided. Full details are listed in Supplementary Table 1.

### Microbial epitope candidate prioritization

Microbial epitopes candidates were prioritized based on their immunogenic potential, employing specific thresholds for epitope-HLA immunogenicity scores, IC50 binding affinities (best and median values, measured in nM), their respective percentile ranks, and microbial abundance. Epitope candidates failing to meet all required thresholds within a given tier were excluded from that tier.

Epitope prioritization was structured into three immunogenicity tiers, defined as follows:

#### Weak immunogenicity

Epitope candidates with an Immunogenicity Score > 0.5, Best and Median IC50 Scores < 500 nM, Best and Median Percentile Ranks < 2, and Microbe Abundance > 0%.

#### Moderate immunogenicity

Epitope candidates with an Immunogenicity Score > 0.6, Best and Median IC50 Scores < 300 nM, Best and Median Percentile Ranks < 1, and Microbe Abundance > 1%.

#### High immunogenicity

Epitope candidates with an Immunogenicity Score > 0.8, Best and Median IC50 Scores < 50 nM, Best and Median Percentile Ranks < 0.5, and Microbe Abundance > 5%.

Since the categorization of epitope-HLA complexes into Highly, Moderately, and Weakly immunogenic tiers may result in an epitope being assigned to more than one category, we adopt the highest tier classification for those epitopes, meaning an epitope categorized as both Highly and Moderately or Weakly will be classified as Highly immunogenic.

### Mutation-derived neoepitope candidate annotation

Comprehensive features and annotations were integrated to support mutation-derived neoepitope candidate evaluation. These included: (i) predicted immunogenicity tier (as determined by MimicNeoAI); (ii)peptide features (mutant and wildtype peptide sequences, peptide length, extended mutant peptides, and their lengths); (iii) HLA allele predicted to present the peptide; (iv)variant annotation (Ensembl gene name and ID, transcript ID, amino acid change, mutation positions, variant classification, transcript biotype, and HGVS nomenclature at both cDNA and protein levels); (v) variant support metrics (tumor DNA depth and variant allele frequency); (vi) expression metrics (TPM expression levels of the gene and transcript encoding the mutant peptide); (vii) population frequency data (allele frequencies in global and ancestry-specific populations); (viii) MHC binding metrics (best and median predicted IC50 values and percentile ranks for mutant and wildtype peptides, fold change, and the corresponding predictive methods); (ix) predicted immunogenicity score; and (x) physicochemical properties (amino acid composition, polarity, volume, net charge, hydrophobicity, Boman index, aliphatic index, and isoelectric point). A comprehensive description of all annotations and features is provided in Supplementary Table 1.

### Mutation-derived neoepitope candidate prioritization

Mutation-derived neoepitope candidates were prioritized using criteria analogous to microbial epitope candidate prioritization, with additional consideration of tumor-specific features. Each candidate was evaluated based on epitope-HLA immunogenicity score, best and median mutant (MT) IC50 values, corresponding percentile ranks, fold-change between mutant and wild-type peptides, and gene expression of the originating gene. Mutation-derived neoepitope candidates failing to meet all specified thresholds within a particular immunogenicity tier were excluded from that tier.

The tiering criteria were defined as follows:

#### Weak immunogenicity

Mutation-derived neoepitope candidates with an Immunogenicity Score > 0.5, Best and Median MT IC50 Scores < 500 nM, Best and Median Fold Changes > 2, Best and Median MT Percentile Ranks < 2, and Gene Expression > 0 TPM.

#### Moderate immunogenicity

Mutation-derived neoepitope candidates with an Immunogenicity Score > 0.6, Best and Median MT IC50 Scores < 300 nM, Best and Median Fold Changes > 4, Best and Median MT Percentile Ranks < 1, and Gene Expression > 1 TPM.

#### High immunogenicity

Mutation-derived neoepitope candidates with an Immunogenicity Score > 0.8, Best and Median MT IC50 Scores < 50 nM, Best and Median Fold Changes > 10, Best and Median MT Percentile Ranks < 0.5, and Gene Expression > 10 TPM.

### Mimicry of tumor neoantigens identification

To identify mimicry of tumor neoantigens, the MimicNeoAI framework performs an exhaustive comparison of all possible pairs between microbial epitope candidates (N=X) and mutation-derived neoepitope candidates (N=Y) using Cartesian product analysis (X×Y pairwise alignments). Each peptide pair is evaluated by analyzing the length of the longest common substring (LCS) normalized by the maximum length of the two peptides. The LCS-based normalization ensures cross-comparability of peptide pairs of varying lengths.

### Transcriptomic analysis

RNA sequencing reads were aligned to GRCh38 using STAR^62^ (v2.5.2b) with parameters --outSAMunmapped Within and --quantMode TranscriptomeSAM. Transcript abundance quantification was subsequently performed using RSEM^63^ (v1.2.28) with the command rsem-calculate-expression --paired-end --alignments.

### ScTCR-seq data analysis and TCR binding probability assessment

Raw scTCR-seq data were obtained in FASTQ format and processed using Cell Ranger VDJ (v7.2.0) with default parameters to identify TCR α- and β-chain sequences for each individual cell, along with corresponding clonotype, sequence, and abundance information. Cells with clonal isoform data were extracted from the scTCR-seq data for further analysis. These cells were subjected to quality control using Fastp (v0.23.2) to remove low-quality reads. Cleaned reads were aligned to the GRCh38 human genome using STAR (v2.7.10a) with default parameters, and UMI count matrices were processed with Seurat^64^ (v4.4.0) for normalization and feature selection. Low-quality cells were excluded from further analysis, and potential doublets were removed using DoubletFinder (v1.10.2). Clustering and dimensionality reduction visualization were performed using Scanpy^65^ (v1.10.2). The first 10 principal components (PCs) were used to construct a k-nearest neighbor (KNN) graph, with each cell connected to its 10 nearest neighbors. The distribution of different TCRs was visualized using UMAP to generate a spatial representation of TCR diversity and clustering patterns. TCR binding probability was predicted using pMTnet^66^ to evaluate the likelihood of TCR-epitope-HLA interactions.

### TCR-epitope-HLA interactions structural modeling

The initial crystal structure of peptide-MHC (pMHC) complex was obtained from the Protein Data Bank (PDB^67^). Non-crystallographic water molecules were removed using PyMOL (v1.7.2.1), and the peptide was separated from the MHC molecule. The target peptide structure was constructed using the mutation tool in PyMOL, based on the original peptide structure. The newly generated target peptide was the docked to the MHC molecule using the HDOCK (v1.1), and the top ten ranked conformations were obtained. Visual analysis in PyMOL was conducted to select the most reasonable binding mode for the pMHC complex model.

For TCR modeling, the variable region amino acid sequences of its α and β chains were submitted to the AlphaFold3 server for de novo prediction. The model with the highest overall confidence was selected for further analysis. This predicted TCR model was then docked to the constructed pMHC complex model using HDOCK (using the pMHC complex as the ligand). Among the multiple predicted models, the final TCR-pMHC complex model was manually selected, considering the docking ranking and the interaction between the TCR β chain CDR3 region and pMHC. This selected model was then used as the initial structure for molecular dynamics simulations.

### TCR-epitope-HLA interactions molecular dynamics simulations

Molecular dynamics simulations were performed using the GROMACS software package and the AMBER99SB-ILDN force field. All bond lengths involving hydrogen atoms were constrained using the LINCS algorithm, enabling a simulation step of 2 fs. Long-range electrostatic interactions were computed using the Particle Mesh Ewald (PME) method, while a cutoff radius of 1.2 nm was applied for short-range electrostatic and van der Waals interactions. The system was placed in a cubic box with a minimum distance of 1.5 nm between the protein complex and the box edge, and was solvated using the SPCE water model. To neutralize the system, appropriate numbers of Na□ or Cl□ ions were added.

Prior to the simulation began, the system underwent energy minimization using the steepest descent method until the maximum force in the system was less than 1000.0 kJ/mol/nm to eliminate unreasonable contacts between atoms. Equilibration was performed in two phases: (i) a 1 ns NVT ensemble (constant temperature and volume) equilibration with position restraints on the protein backbone, maintaining the system temperature at 300 K using a V-rescale thermostat; followed by (ii) a 1 ns NPT ensemble (constant temperature and pressure) equilibration under position restraints, controlling the system pressure at 1.0 bar while maintaining the temperature at 300 K using a Parrinello-Rahman barostat. After equilibration, all position restraints were removed, and a 100 ns production molecular dynamic simulation was conducted.

To compute the binding free energy between TCR and pMHC, the trajectory from the production simulation was first processed using the gmx trjconv tool (with the -pbc mol and -center parameters) to corrected for periodic boundary conditions and ensure the complex remained intact and centered within the simulation box in each frame. The resulting trajectory was then used as input for the gmx_MMPBSA tool, applying the generalized Born (GB) model (igb=5) to calculate the TCR-pMHC binding free energy.

### Molecular dynamics simulations setup and computational configuration

The aforementioned MD simulations were performed on a server equipped with an Intel(R) Xeon(R) Gold 6342 CPU @ 2.80GHz (24 cores) and two 3090 GPUs, each with 24GB of memory, utilizing GPU acceleration for computation. Each simulation of TCR-epitope-HLA interactions was accelerated using 12 CPU cores and one 3090 GPU. The simulation times varied between 24 to 48 hours. A total of three nodes were used for parallel computation.

## Supporting information

Features considered by MimicNeoAI

Epitope-HLA complexes used for model construction and performance

Performance comparison of MimicNeoAI and other algorithms using experimental validated microbial epitopes

Microbial epitope and mutation-derived neoepitope candidates identified by MimicNeoAI

Mimicry of tumor neoantigen pairs detected from CRC sample

Predicted binding probability of microbial epitope-HLAs complex with TCRs

## Code availability

MimicNeoAI is packaged, and distributed as an open-source, publicly available repository at https://github.com/BioStaCs-public/MimicNeoAI

## Data availability

The raw sequencing data generated in this study are available in the SRA under accession number SUB15514373*(controlled access; contact corresponding author for authorization)*. The trained model, custom microbial reference database, as well as the packed pipeline code, are all available on Zenodo at 10.5281/zenodo.15582924.

## Acknowledgements

We sincerely express our gratitude to the Gastrointestinal oncology Integrated Research Team of Hefei Cancer Hospital of CAS for their assistance in sample extraction, data organization, standardization of clinical cases and pathological prognosis evaluation. We gratefully acknowledge the Hefei Advanced Computing Center and Virtaitech (Shanghai) Co., Ltd for providing computational resources and technical support. We thank the colleagues in our research group for inspiring discussion and their contributions.

## Author contributions

Y.Z., T.C., J.F., and L.Z conceptualized the study. T.C. was responsible for the model design and pipeline implementation. T.C., Y.Z., M.L., W.W and X.Z. performed data analysis and model evaluation. Z.L., Y.H., Y.Z., F.Y. B.Z Q. J and H.L. provided clinical samples. T.C., W.W. and Y.Z. wrote the manuscript. L.Z., J.F., and Y.Z. supervised the study.

## Supplementary

### Supplementary information

#### Generation of epitope candidates

Central to the MimicNeoAI framework, the microbial epitope candidate detection sub-pipeline is adept at identifying low-abundance tumor-associated microbial epitope candidates from WTS data. This methodology enables the detection of microbial epitope candidates even in low-biomass environments, with compatibility for large public datasets facilitating scalable, pan-cancer analyses. Initially, cleaned sequencing reads undergo processing to eliminate host-derived sequences, after which the remaining non-host reads are aligned to a custom-built microbial genome reference, minimizing contamination and enhancing microbiome identification. This reference is constructed through a combination of LLM-based annotation and manual curation, ensuring the inclusion of microorganisms capable of colonizing the human body or influencing cancer progression or anti-tumor treatment response. Subsequently, microbial reads were translated into proteins and matched against protein databases. A rigorous filtering process is applied of the alignment results, allowing for the precise identification of microbial peptides that may serve as epitope candidates, thereby enhancing the overall accuracy of epitope detection.

The mutation-derived epitope candidate detection sub-pipeline of MimicNeoAI is designed to detect mutation-derived epitope from WES data. This process begins with the alignment of cleaned sequencing reads to the human reference genome, identifying tumor-specific somatic mutations that are subsequently filtered to exclude low-quality variants. Additionally, expression quantification from WTS data is utilized, retaining only mutations in genes with detectable expression, thereby ensuring that identified mutations are actively transcribed and more likely to be presented as mutation-derived epitopes.

#### Prediction and prioritization of epitope immunogenicity

To enhance the prioritization of peptides with significant immunogenic potential, MimicNeoAI incorporates HLA typing from WTS or WES data. Building on the peptide candidates generated by the aforementioned sub-pipelines, the framework transcends basic peptide-HLA binding probability predictions. Utilizing a variety of algorithms to calculate binding affinities, MimicNeoAI further integrates an immunogenicity prediction model. This model leverages BiLSTM networks to interpret long-range amino acid interactions within peptide-HLA complexes, complemented by a Fully Connected Neural Network (FCNN) that assessing eight physicochemical properties (Amino acid composition, polarity, residue volume, net charge, hydrophobicity, Boman index, aliphatic index, and isoelectric point).

During the post-model screening steps, MimicNeoAI employs an extensive selection and ranking process, considering characteristics of both microbial and host-derived peptides generated from the aforementioned pipeline. The microbial peptides characteristics include metrics such as total library size, microbial reads ratio, and microbial abundance derived from nucleic acid-level microbial alignment, along with protein reference sequence alignment results (e.g., percentage identity, gap opening, e-value, and bitscore). In contrast, the host-derived peptides characteristics encompass DNA-level metrics (e.g., depth, variant allele frequency, and allele frequency), gene expression levels, mutation details (positions, quality scores), and protein-level metrics (split protein positions). By integrating binding probability and immunogenicity scores, MimicNeoAI effectively ranks the microbial and mutation-derived epitope candidates, categorizing them into non-immunogenic, weakly immunogenic, moderately immunogenic, and highly immunogenic classes.

#### Materials of immunogenicity prediction model

The immunogenicity prediction model of MimicNeoAI was trained using peptide-HLA complexes obtained from multiple sources (Supplementary Table 2). The host-derived (human) dataset, compiled from the IEDB, includes 117 HLA-I alleles and 12,039 peptides, resulting in 15,482 peptide-HLA complexes, primarily sourced through two specialized algorithms: DeepHLApan and DeepImmuno. This dataset was balanced with 7,741 immunogenic and 7,741 non-immunogenic complexes. The microbial dataset, derived from MicroEpitope, includes 4,556 HLA-I and 650 HLA-II alleles, with 17,894 HLA-I-binding and 1,018 HLA-II-binding peptides. These yielded 28,246 peptide-HLA-I complexes (14,123 immunogenic, 14,123 synthetic non-immunogenic) and 1,754 peptide-HLA-II complexes (877 immunogenic, 877 synthetic non-immunogenic), ensuring balanced positive/negative representation, and the non-immunogenic complexes were randomly generated. Additionally, we compiled a microbial epitope validation set of 46 peptide-HLA complexes. To ensure independence, we excluded any peptides present in the training set, leaving 42 peptide-HLA complexes for validation. This set combines 21 literature-verified immunogenic microbial complexes and 21 randomly generated non-immunogenic complexes for unbiased evaluation.

## Supplementary figures

**Supplementary Figure 1.**
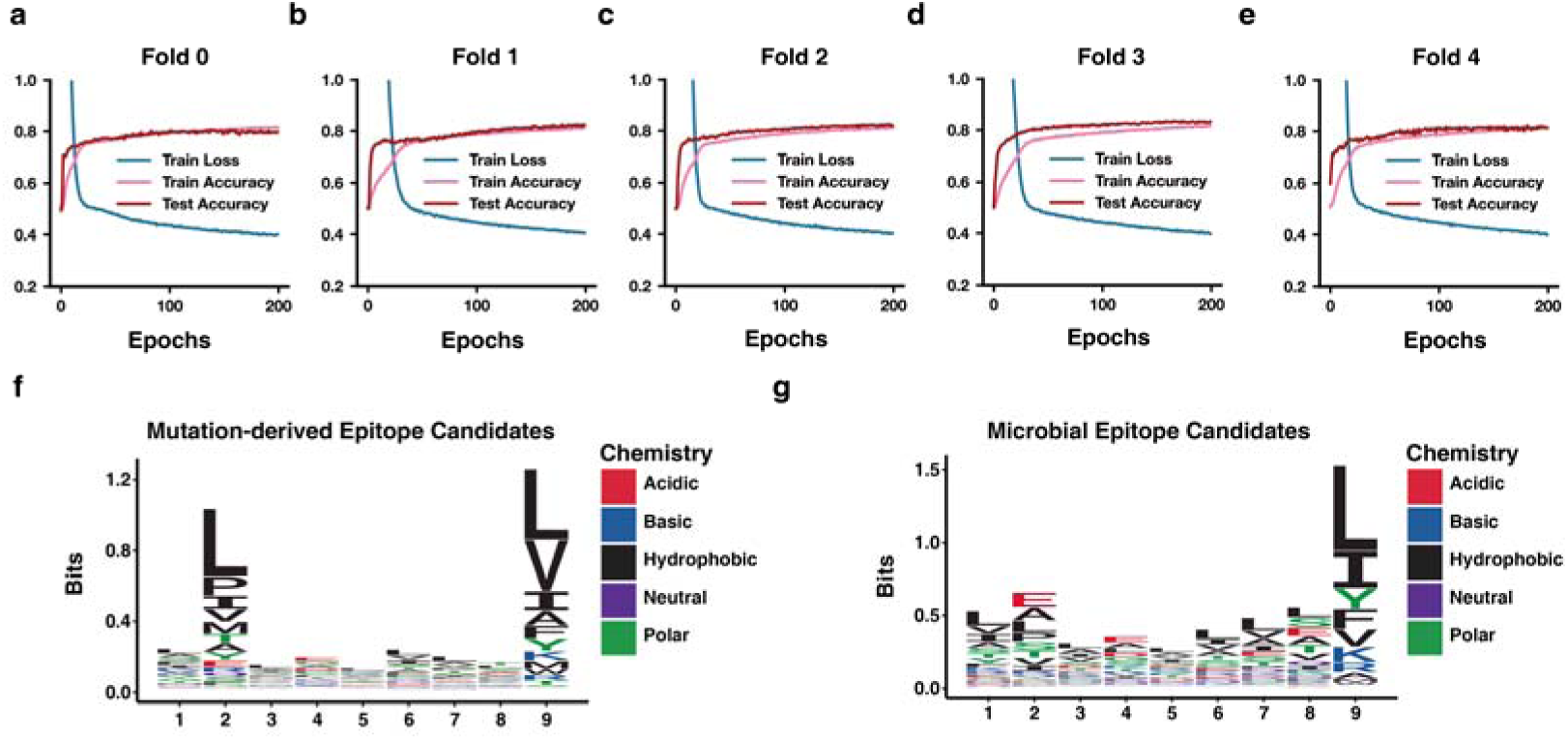
Evaluation of training performance via five-fold cross-validation and amino acid sequence features. (**a-e**) Plots of train loss (blue), train accuracy (pink), and test accuracy (red) for each fold of the five-fold cross-validation. Sequence logo plots of immunogenic peptides with length 9 from the host-derived dataset (**f**) and microbial dataset (**g**), showing the amino acid frequency distribution in positive training samples.

**Supplementary Figure 2.**
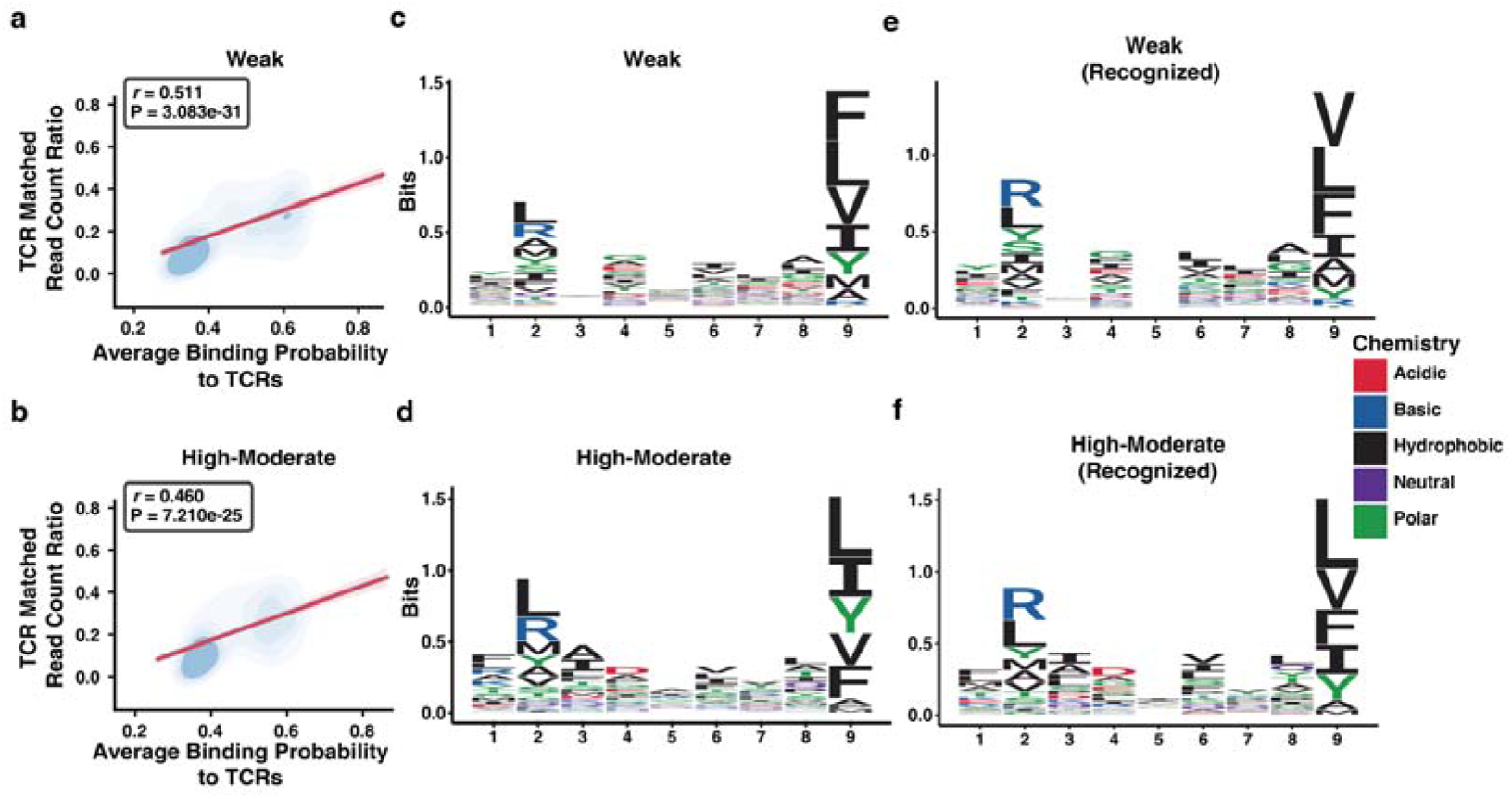
Analysis of T cell recognition and amino acid frequency in microbial epitope candidates. (**a)** Correlation between T cell read count ratios and average binding probability to TCRs in the weak immunogenicity group of microbial epitope candidates. (**b**) Correlation between TCR read count ratios and average binding probability to TCRs in the high-moderate immunogenicity group of microbial epitope candidates. Sequence logo plots of microbial epitope candidates with length 9 in the weak (**c**) and High-Moderate group (**d**). Sequence logo plots of recognized microbial epitope candidates with length 9 in the weak (**e**) and high-moderate immunogenicity group (**f**).

**Supplementary Figure 3.**
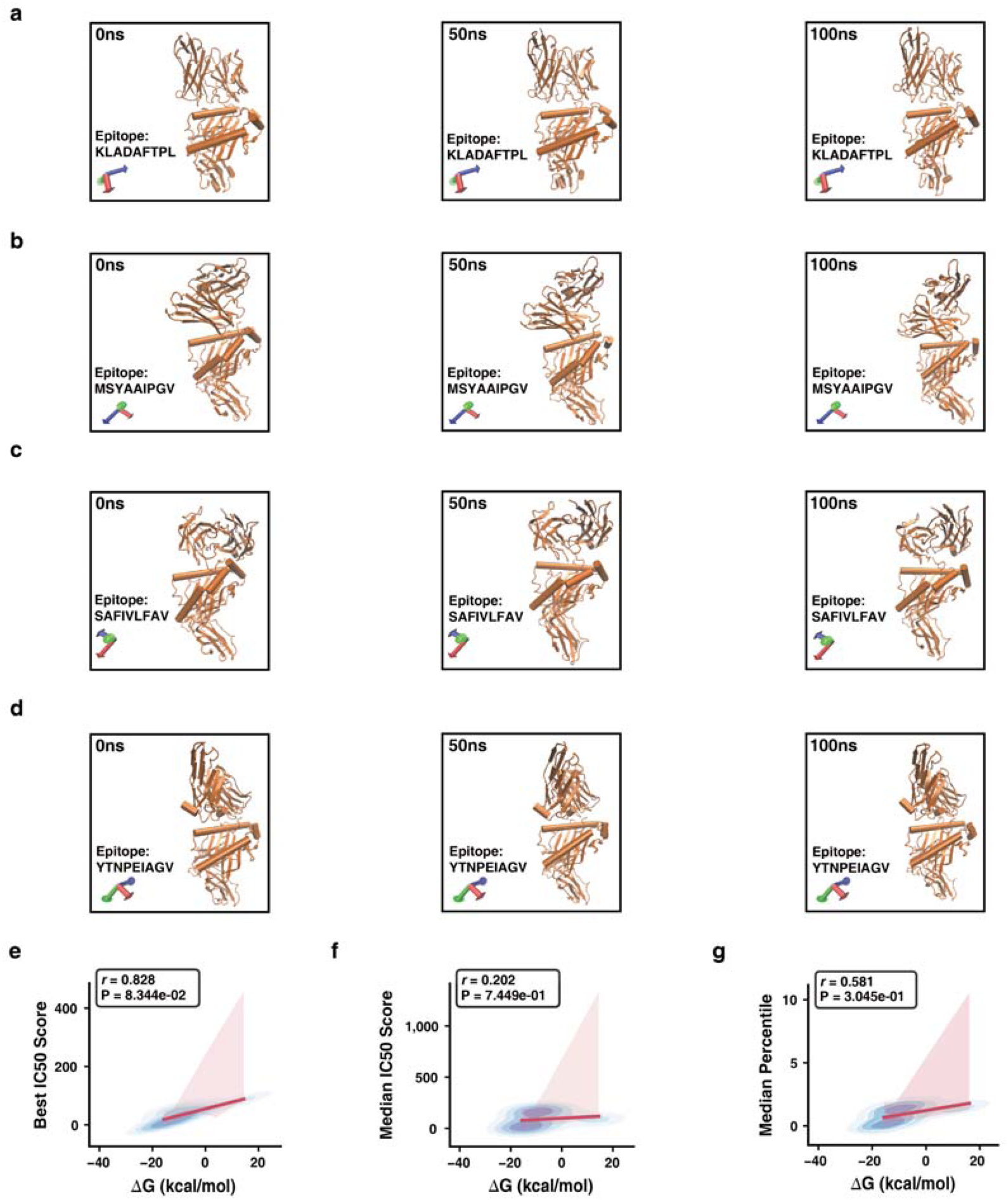
Evaluation of HLA binding probability characteristics through pearson correlation analysis and validation of TCR-epitope-HLA trajectory stability. (**a-d**) Molecular dynamics simulation trajectory of Clonotype2 and HLA-A02:01 in complex with the epitope candidates KLADAFTPL (**a**), MSYAAIPGV (**b**), SAFIVLFAV (**c**), and YTNPEIAGV (**d**) is divided into three critical time frames (0 ns, 50 ns, and 100 ns). (**e)** The correlation between the best IC50 Scores of microbial epitope candidates and their ΔG. (**f)** Pearson correlation between the median IC50 scores and ΔG. (**g)** Pearson correlation between the median percentiles of microbial epitope candidates and their ΔG.

